# Early Identification of At-Risk Patients: Proteomic Signature Predicts Progression to Decompensated Cirrhosis

**DOI:** 10.64898/2026.03.04.709475

**Authors:** Marie Louise N. Therkelsen, Nicolai J. Wewer Albrechtsen, Mikkel Parsberg Werge, Mira Thing, Puria Nabilou, Elias Badal Rashu, Liv Eline Hetland, Signe Boye Knudsen, Anders Ellekær Junker, Elisabeth Douglas Galsgaard, Jesper Velgaard Olsen, Mads Grønborg, Nina Kimer, Lise Lotte Gluud

**Author notes:** **Corresponding Author:** Lise Lotte Gluud, Gastro Unit, Copenhagen University Hospital Hvidovre, Kettegaard Alle 30, 2650 Hvidovre, Denmark.

## Abstract

**Background & Aims:** Early identification of decompensation in patients with cirrhosis is important to enable timely detection, management of complications and for effective treatment. This study investigates the biology of decompensation and aim to identify protein biomarkers for identification of high-risk patients.

**Methods:** The primary analysis included plasma samples from 46 patients with metabolic dysfunction associated steatotic liver disease (MASLD) related cirrhosis. Plasma samples were depleted for the top 14 most abundant proteins and the proteome was measured by liquid chromatography tandem mass spectrometry. The dataset was divided into a training (14 compensated, 10 decompensated) and a test cohort of compensated patients (11 progressing to decompensation, 11 remaining compensated). Changes in protein levels were determined by ANCOVA and a prognostic model was developed using logistic regression. External validation was performed in an independent cohort of 120 patients with alcohol-related cirrhosis. Time-to-event analyses were conducted in this cohort using Cox regression.

**Results:** 52 proteins involved in impaired hepatic function, fibrogenesis, immune activation, and metabolic changes were significantly different between compensated and decompensated patients. A prognostic model with four proteins (NBL1, LTBP4, APOC4, GHR), demonstrated predictive ability for future decompensation (AUC=0.93, 73% sensitivity, 100% specificity). In the external validation cohort, the model demonstrated generalizability (AUC=0.78, 72% sensitivity, 82% specificity). Validation cohort time-to-event analyses showed that higher baseline scores were associated with shorter time to liver-related events (HR 1.32; log-rank *p* = 0.027), underscoring the panel’s prognostic value.

**Conclusion:** Our study indicates that patients with decompensated cirrhosis are characterized by proteomic signatures of fibrogenesis and metabolic dysfunction. Capturing these signatures could help identify patients at risk of complications and potentially those eligible for aetiology directed treatment.

**Impact and Implications:** Addressing a critical unmet need for early detection of cirrhosis decompensation, our proteomic study identifies a four-protein panel with predictive ability for decompensation. These findings hold significant implications for hepatologists, clinical researchers, and healthcare systems, offering a novel tool to enhance prognostication and refine treatment strategies, potentially facilitating targeted patient monitoring. However, considering the small discovery sample size and the distinct aetiology of the external validation cohort, further validation is essential before broad clinical integration.

**Graphical Abstract:** 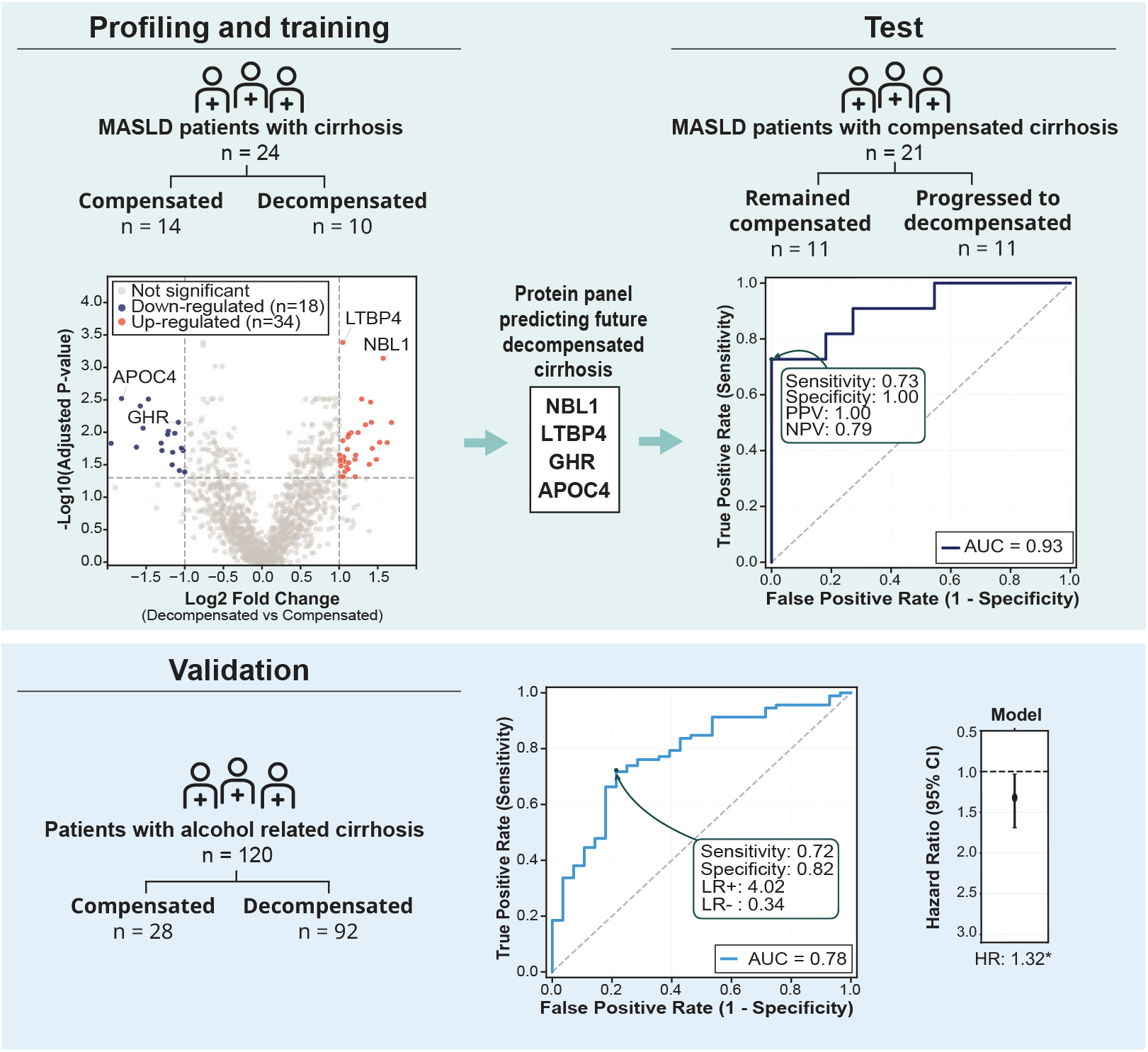

## Introduction

Metabolic dysfunction-associated steatohepatitis (MASH) poses a significant global health challenge [2]. Without timely intervention, MASH can progress to advanced fibrosis [3] and ultimately to decompensated cirrhosis, with complications such as ascites, hepatic encephalopathy (HE), and variceal bleeding [4]. This progression not only marks a severe worsening of the patient’s condition and a significantly poorer prognosis but also notably increases the risk of HCC [5]. Early identification of patients at risk of progressing from compensated to decompensated cirrhosis is crucial, as it enables intensified monitoring and timely, aetiology-directed interventions to prevent complications and improve patient outcomes [6, 7]. Current clinical tools for predicting decompensation remain limited, and there is an unmet need for biomarkers that can accurately stratify risk in patients with compensated cirrhosis.

In this study, we utilize plasma proteomics to identify biomarkers capable of distinguishing patients at risk of decompensation from those who remain compensated. We perform proteome profiling on samples from patients with Metabolic dysfunction-Associated Steatotic Liver Disease (MASLD) cirrhosis with subsequent validation and assessment of generalizability in an independent cohort including patients with alcohol-related cirrhosis. Our aim is to identify and validate a protein-based prognostic signature for early identification of high-risk patients who may benefit from closer surveillance and targeted therapeutic interventions.

## Materials & Methods

### Primary Study Population

This study included participants with histologically verified compensated cirrhosis from a prospective cohort recruiting patients from the outpatient clinic at the Gastro Unit, Copenhagen University Hospital Hvidovre (Copenhagen Cohort of MASLD, Diabetes and Obesity (COCOM-DO)). Ethical approval was obtained from the Research Ethics Committee of the Capital Region of Denmark (H-17029039), and the study was conducted in accordance with the 1975 Declaration of Helsinki. Following written consent, participant who fulfilled the diagnostic criteria for MASLD underwent clinical assessment, blood tests, Fibrosis-4 Index (FIB-4), liver stiffness measurement (LSM), and a liver biopsy if significant or advanced fibrosis was suspected (described in [8]). Patients were subsequently followed yearly with clinical assessments, blood tests, and LSM. Plasma samples were collected at baseline and subsequently once yearly. All participants were fasting at the time of collection. Venous blood samples were collected into EDTA tubes and centrifuged at 2500xg for 10 minutes at 4°C. Plasma was then separated and stored at −80°C until the time of analysis. This study analyzed plasma samples from 46 patients included from 2019 to 2023 with biopsy-confirmed cirrhosis. Decompensation (development of ascites, jaundice, hepatic encephalopathy, or variceal bleeding), was assessed in August 2024.

### Independent Validation Cohort

To validate the findings and to assess the generalizability beyond MASH aetiology, we performed external validation on an independent cohort of patients with alcohol-related cirrhosis (the PROspective cohort study of Disease and Outcomes in Cirrhosis (PRO-DOC2019), the Gastro Unit, Copenhagen University Hospital Hvidovre). Ethical approval was obtained from the Research Ethics Committee of the Capital Region of Denmark (H-19024348). The cohort included 28 patients with compensated cirrhosis and 92 with decompensated cirrhosis. Baseline plasma samples were collected as the COCOM-DO samples.

### LC-MS/MS analysis

An automated plasma proteomics sample preparation platform was employed using a Tecan Fluent liquid-handling robot. Highly abundant plasma proteins were depleted from 5 *µ*L plasma using Top14 Abundant Protein Depletion Resin (Thermo Scientific, cat# A36372). The remaining proteins were digested in-solution into peptides with Trypsin/Lys-C (Promega, cat# V5071). Following digestion, peptides were acidified and loaded onto Evotips (EvoSep Biosystems, cat# EV2013) according to the manufacturer’s protocol. liquid chromatography-tandem mass spectrometry (LC-MS/MS) analysis was performed using an Orbitrap Astral (Thermo Scientific) coupled to an Evosep One (EvoSep Biosystems). Samples were analyzed via single shot narrow-window data-independent acquisition (nDIA) with an 11-minute gradient on an 8cm Evosep column (EV1109, EvoSep). Raw LC-MS files were processed using Spectronaut v18 (Biognosys). DIA-MS/MS spectra were searched using the spectral library-free direct-DIA algorithm against a human UniProt FASTA database (version 20250203, 20,433 entries). On average, 1766 proteins were identified per sample. In total 1529 proteins were quantified in at least 70% of runs, while 753 proteins were quantified across all samples (100% completeness). Protein intensities were *log*_2_-transformed. Quantile normalization and batch correction for plate effects were applied using the limma R package. After preprocessing, protein groups were retained only if at least 70% of samples had valid (non-missing) values.

### Statistical Analysis

The primary data set was divided into a training cohort (n=24; 10 decompensated and 14 compensated) and a test cohort (n=22) comprising patients with compensated cirrhosis at baseline: 11 who later developed decompensation and 11 who remained compensated. The split in compensated patients was selected at random (with a fixed seed). Analyses were conducted in Python (version 3.11.9) using statsmodels, scikit-learn, matplotlib, seaborn and lifelines. Baseline characteristics are presented as median and SD for normally distributed continuous variables, as the median and interquartile range (IQR) for non-normally distributed continuous variables, and as counts for discrete data. Normality was assessed using the Shapiro-Wilk test. Group differences were assessed using either ANOVA, the Wilcoxon test, Student’s T-test, or the Chi-square test. Differential protein abundance was evaluated in the training cohort using analysis of covariance (ANCOVA) adjusting for age, sex, and type 2 diabetes (T2D), and corrected for multiple testing using the Benjamini–Hochberg false discovery rate (FDR). Results shown as volcano plots, with log2 fold change and −*log*_10_(FDR-adjusted p-value). Statistical significance was defined as an FDR-adjusted p-value ≤ 0.05, and a fold change ≥ 2. Heatmap was generated with regulated proteins (protein-wise z-scores). To select a protein panel, we performed feature selection in the training cohort using an L1-penalized logistic regression (LASSO). We started with differentially regulated proteins (n=52), standardized using z-score normalization, and included clinical covariates (age, sex). Model tuning and evaluation were conducted via 5-fold cross-validation, optimizing AUC. The regularization strength (C, inverse of *λ*) was searched from 10^−3^ to 10^2^. The final selection comprised all proteins with non-zero coefficients after regularization. Given the sample size (n=24), we restricted the panel to the top four proteins. These proteins were used to fit a final model using L2-penalized logistic regression (Ridge regression, C=0.1) to minimize overfitting. The final model was evaluated on the test cohort (n=22) to assess its ability to identify compensated patients who would later progress to decompensated cirrhosis. Model performance was assessed using ROC curves, with AUC and 95% CI estimated by bootstrap resampling (2000 iterations). Youden’s threshold (sensitivity + specificity − 1) was evaluated, with corresponding sensitivity, specificity, positive predictive value (PPV), negative predictive value (NPV), and positive and negative likelihood ratios (LR) and confusion matrices.

For the validation cohort, model predictions were generated using the fitted model without re-tuning. Cox proportional hazards regression was used to analyze time to liver-related events (admission with decompensation after baseline or liver-related death). Two approaches were employed: first, model predictions were logit-transformed to normalize the probability distribution, with hazard ratio (HR) reported per unit increase in log odds; second, individual protein intensities were standardized, with HRs reported per 1-SD increase. Time was measured in days from baseline visit to the first event or censoring at the end of study (median 1120 days). P-values were FDR-corrected (Benjamini-Hochberg), and significance levels indicated (\**p <* 0.05, \*\**p <* 0.01, \*\*\**p <* 0.001). Kaplan-Meier illustrated event-free stratified by Youden’s threshold for model-based analysis and median intensity for proteins.

## Results

The median age of participants in the primary analysis was 62 years, 61% were female and 70% had T2D. Baseline characteristics were similar between groups in both training and testing datasets (Table 1). As expected, platelet counts and albumin were significantly lower in the decompensated group across both datasets.

**Table 1:** Baseline characteristics. Groups in the training and testing datasets. Continuous variables are reported as median (SD) for normally distributed data. Categorical variables are reported as counts (percentages). Differences between groups was assessed using Student’s t-test or the Chi-square test. IHD = Ischemic heart disease.

|  | Training |  |  | Testing |  |  |
| --- | --- | --- | --- | --- | --- | --- |
| | Compensated | Decompensated | $p$ | Compensated | Decompensated later | $p$ |
| **Number** | 14 | 10 | — | 11 | 11 | — |
| **Age**, median (SD) | 62 (9.9) | 67 (8.0) | 0.181 | 58 (11.2) | 61 (11.4) | 0.481 |
| **BMI**, median (SD) | 32.6 (5.0) | 30.1 (4.1) | 0.353 | 33.5 (5.5) | 30.5 (4.9) | 0.353 |
| **Female**, n (%) | 6 (43%) | 8 (80%) | 0.162 | 6 (55%) | 8 (73%) | 0.658 |
| **Type 2 diabetes**, n (%) | 10 (71%) | 9 (90%) | 0.552 | 6 (55%) | 7 (64%) | 1 |
| **Hypertension**, n (%) | 10 (71%) | 6 (60%) | 0.884 | 5 (46%) | 8 (73%) | 0.385 |
| **Dyslipidemia**, n (%) | 11 (79%) | 7 (70%) | 1.0 | 6 (55%) | 7 (64%) | 1 |
| **IHD**, n (%) | 1 (7%) | 3 (30.0) | 0.354 | 1 (9.1) | 0 (0.0) | 1 |
| **Platelets**, median (SD) | 191 (47) | 135 (50.3) | 0.047 | 202 (41.6) | 109 (38.7) | 0.001 |
| **Albumin**, median (SD) | 39.5 (4.1) | 33.5 (5.3) | 0.002 | 40.0 (3.2) | 31.0 (5.0) | 0.001 |
| **Triglycerides**, median (SD) | 2.3 (0.9) | 1.3 (0.6) | 0.005 | 1.9 (0.7) | 1.6 (0.7) | 0.511 |

### Protein profile were significantly different between patients with decompensated versus compensated cirrhosis

We identified 52 plasma proteins that were significantly different between patients with decompensated and compensated cirrhosis (adjusted p-value ≤ 0.05, fold change ≥ 2), comprising 18 downregulated and 34 upregulated proteins (Figure 1).

**Fig. 1:**
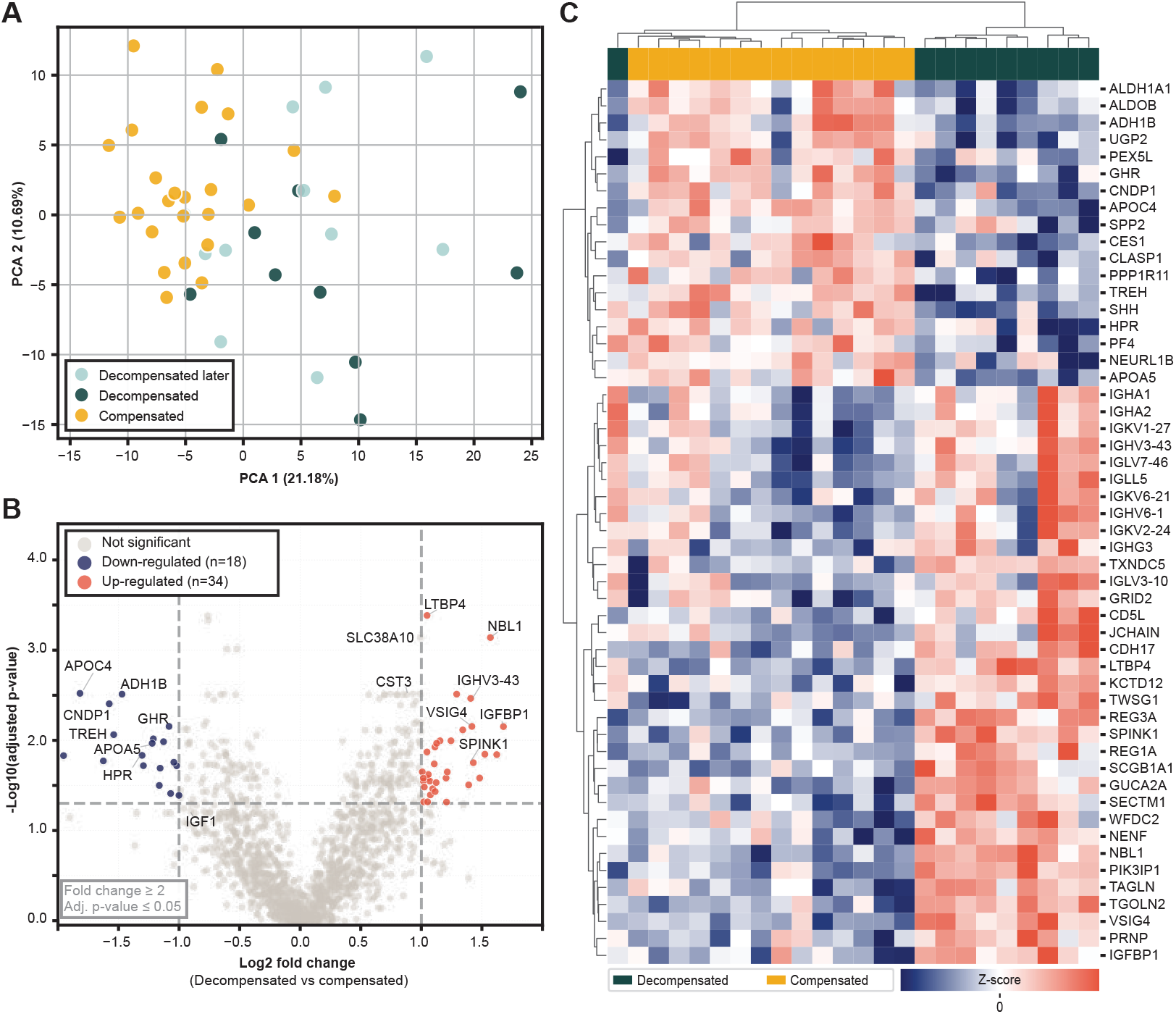
Protein regulation. (A) principal component analysis (PCA) of included patients based on protein intensities (100% valid values). Colored by group. (B) Volcano plot of differential protein intensities in decompensated versus compensated cirrhosis (log_2_ fold change and −log_10_ adjusted p-value). Significantly down-regulated (blue) and up-regulated (red). Dashed lines indicate significance cutoffs (fold change ≥ 2 and adjusted p-value ≤ 0.05). (C) Heatmap of significantly regulated proteins (n=52). Values are z-scored (color bar); rows are proteins (gene symbols) and columns are samples. Hierarchical clustering of proteins and samples. Columns are color-coded by group). Training dataset.

The protein profile indicated impaired hepatic function: downregulation of apolipoproteins APOA5 and APOC4 could indicate disrupted hepatic lipid metabolism; reduced growth hormone receptor (GHR) and insulin-like growth factor 1 (IGF1), together with increased insulin-like growth factor-binding protein 1 (IGFBP1), pointed to dysregulation of the GH–IGF1 axis; and low levels of hepatic enzymes (fructose-bisphosphate aldolase B (ALDOB), all-trans-retinol dehydrogenase (ADH1B1), aldehyde dehydrogenase 1A1 (ALDH1A1)) further reflected a decline in metabolic capacity. In parallel, upregulation of latent TGF-*β*–binding protein 4 (LTBP4), neuroblastoma suppressor of tumorigenicity 1 (NBL1), and twisted gastrulation protein homolog 1 (TWSG1) suggested progressive fibrogenesis. Immune activation was also evident, with increased immunoglobulin components (IGHA1, IGHG3, IGKV1-27, IGHV6-1, IGHV3-43, JCHAIN), indicating altered innate immune regulation. Signals of systemic manifestations were evident with upregulation of cystatin C (CST3), associated with renal dysfunction [9], and solute carrier family 38 member 10 (SLC38A10), a sodium-coupled neutral amino acid transporter implicated in the glutamate–glutamine cycle, potentially contributing to HE. Additionally, several proteins had established associations with HCC: carnosine dipeptidase 1 (CNDP1) [10], LTBP4 [11], JCHAIN [12], and ADH1B1 [13].

### Predictive model for liver decompensation in cirrhosis patients

Feature selection on proteins with plasma levels significantly different between groups (n=52), identified 13 proteins. The top four proteins; LTBP4, NBL1, GHR, and APOC4, were selected for the final regression model (Figure 2.A). In the test set (n=22), the model demonstrated strong discriminative ability for predicting future decompensation in initially compensated patients (AUC=0.93, 95% CI: 0.88-1.00). With a Youden’s threshold of 0.49, the model demonstrated robust performance with 100% specificity and PPV (likelihood ratio (LR)+=inf), ensuring that all patients who did not develop decompensation were correctly identified, while it successfully detected the majority of those who later progressed to decompensated cirrhosis (73% sensitivity, 79% NPV, LR-=0.27) (Figure 2.B).

**Fig. 2:**
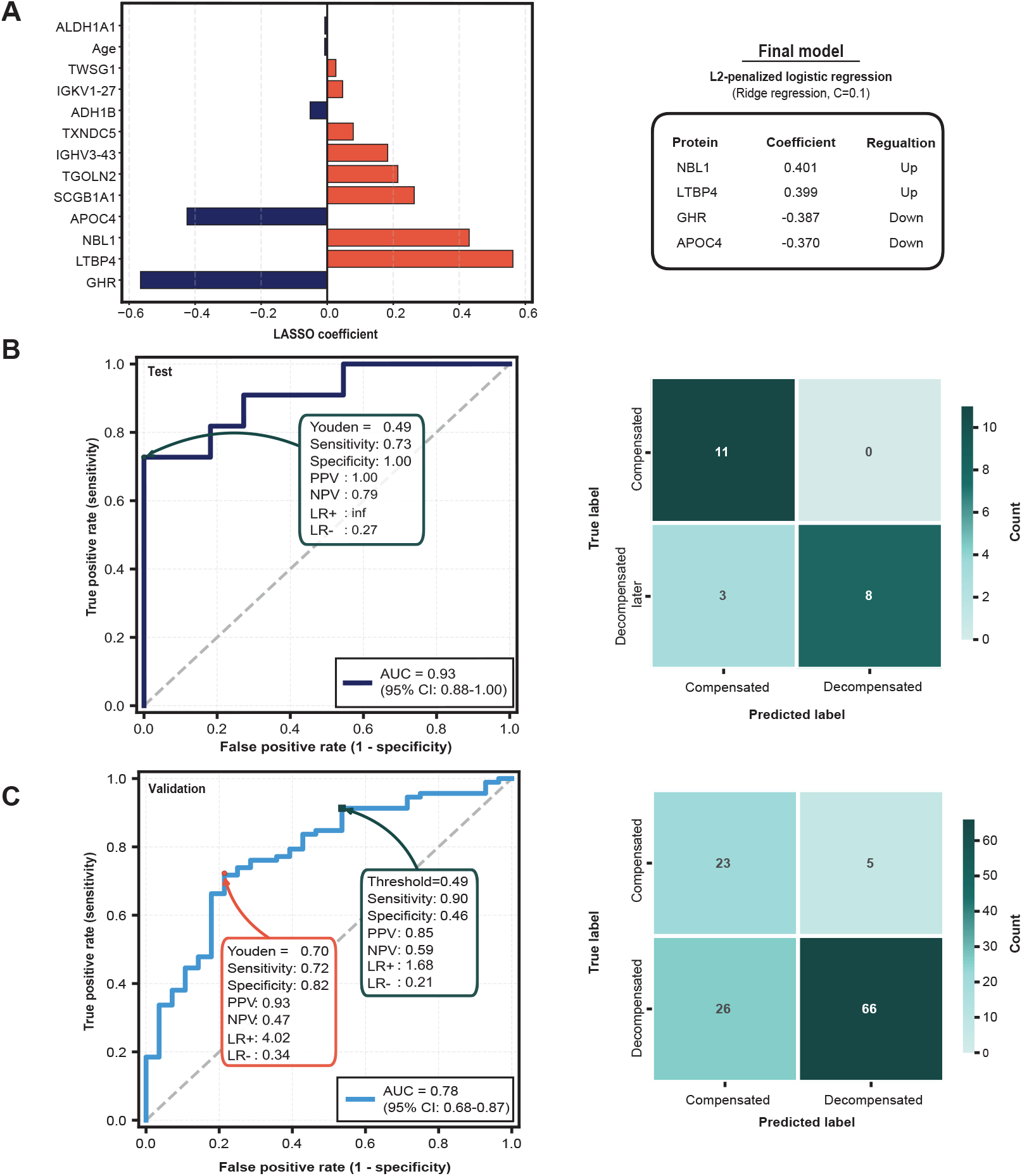
Feature selection and model performance. (A) LASSO regression coefficients for proteins with non-zero coefficients after regularization (L1-penalized logistic regression). Right, selected panel and coefficients (regularization strength C, inverse of *λ*). (B) Model performance on test set (n=22). ROC curves display AUC with 95% CI. Optimal threshold by Youden^*′*^s threshold with corresponding sensitivity, specificity, positive (PPV) and negative (NPV) predictive values, and positive (LR+) and negative (LR-) likelihood ratios. Right, confusion matrix constructed using the optimal threshold. (C) Model performance on validation set (n=120), Two thresholds: validation-specific Youden^*′*^s threshold and the test set’s optimal threshold. Confusion matrix using validation-specific Youden threshold.

**Fig. 3:**
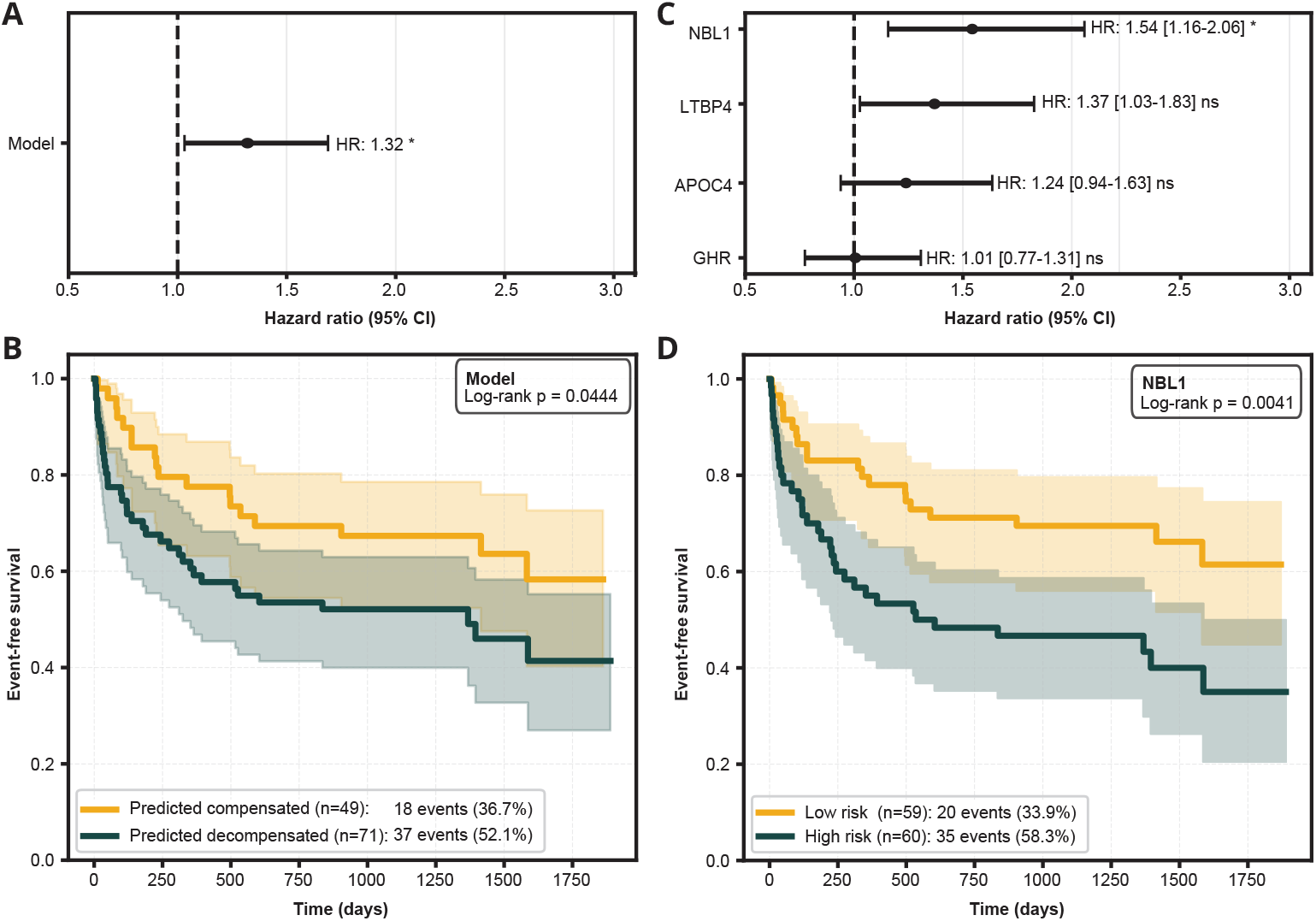
Prognostic value of model and individual proteins in the validation cohort. (A) Forest plot showing hazard ratio (HR) with 95% CI for predicting liver-related events (Cox regression) using the models probability score. (B) Kaplan-Meier curves stratified by model prediction threshold (0.70). (C) Forest plot for individual panel protein (standardized intensities). (D) Kaplan-Meier curves for NBL1, stratified by median NBL1 intensity. FDR-corrected p-values (Benjamini-Hochberg). \**p <* 0.05, not significant (ns).

In the external validation cohort (n=120, alcohol-related cirrhosis), the model exhibited discriminative ability (AUC=0.78) (Figure 2.C). When the test set optimal threshold of 0.49 was applied, the model demonstrated a sensitivity of 90% and a specificity of 46%, with LR+ of 1.68 and LR-of 0.21. In contrast, at the validation cohort’s Youden’s threshold of 0.70, the model achieved a sensitivity of 72% and a specificity of 82%, with LR+ of 4.02 and LR-of 0.34. In the validation cohort, time-to-event analysis showed the model score predicted liver-related events (HR=1.32, 95% CI 1.03–1.69; p=0.027; Figure 3.A). Using the 0.70 threshold, Kaplan-Meier analysis showed significant risk stratification (log-rank *p* = 0.0444; Figure 3.B), with high-risk patients experiencing 52.1% events vs 36.7% in the low-risk group. We further evaluated the individual prognostic value of the four proteins comprising the model (Figure 3.C). Only NBL1 showed significant prognostic value (HR = 1.54, 95% CI: 1.03–1.83; *p* = 0.013) with significant risk stratification (log-rank *p* = 0.0041; Figure 3.D).

## Discussion

The progression from compensated to decompensated cirrhosis represents a pivotal and high-risk transition for patients. Predicting this transition remains a significant clinical challenge, despite its profound implications for prognosis and management. In this study, we addressed this critical need by employing a two-stage approach leveraging plasma proteomics from longitudinal samples. Our methodology aimed to investigate the key pathways affected prior to decompensation and identify biomarkers capable of distinguishing patients approaching decompensation from those who remain stable.

We identified 52 differentially regulated proteins between compensated and decompensated cirrhosis, highlighting the biological shifts accompanying decompensation, including impaired liver function, intensified fibrogenesis, immune dysregulation, and systemic complications. Markers of impaired hepatic lipoprotein synthesis were prominent, with downregulation of APOA5 and APOC4 correlating with lower triglyceride levels. While decreased triglycerides are a known feature of cirrhosis [14], our data further revealed that this decline intensifies with decompensation. Furthermore, the GH–IGF1 axis showed clear dysregulation, evidenced by reduced GHR and IGF1 alongside increased IGFBP1. Impaired insulin signalling, common in cirrhosis, may reduce GHR levels as insulin helps maintain its expression [15]. This consequently diminishes IGF1, given GHR’s crucial role in mediating growth hormone (GH) signalling for hepatic IGF1 production. Altered insulin levels likely elevate IGFBP1, which, as a binding protein to IGF1, further diminishes its activity [16, 15]. Consequently, low circulating IGF1 serves as a critical indicator of impaired hepatic metabolic homeostasis and is associated with worse patient outcomes [17, 18]. Downregulated liver enzymes further indicate compromised hepatocyte metabolic competence: reduced ALDOB suggests impaired glycolytic/fructose-catabolic flux, while decreased ADH1B/ALDH1A1 signify defects in aldehyde detoxification and retinoic acid biosynthesis, pathways widely dysregulated in cirrhosis [13]. The protein profile further indicated progressive fibrogenesis through an activated bone morphometric proteins (BMP) pathway with upregulation of NBL1 and TWSG1, both BMP modulators [19, 20, 21]. Intensified immune regulation was also prominent, the increased immunoglobulin components signify a robust, antigen-driven humoral immune response, while elevated V-set and immunoglobulin domain-containing protein 4 (VSIG4), a complement receptor on tissue-resident macrophages, is known to contribute to immune homeostasis by dampening T-cell responses and inflammation. However, its upregulation in advanced disease can paradoxically facilitate tumor immune evasion [22]. The dysregulation of proteins with established links to HCC (CNDP1, JCHAIN, LTBP4, and ADH1B1) further underscores the malignant potential in cirrhosis progression [10, 12, 11, 13]. Finally, the upregulation of systemic indicators, such as SLC38A10 (might be involved in HE) and CST3 (reflecting renal dysfunction), provides crucial insight into the systemic derangements that are already in motion, often preceding the clinical manifestation of decompensated cirrhosis [9]. Collectively, these molecular alterations, spanning metabolic, fibrotic, immune, and systemic pathways, underscore the severe, multi-faceted, and progressive nature of the disease.

Leveraging these comprehensive molecular insights, we aimed to develop a clinically relevant tool for patient stratification and the early prediction of hepatic decompensation. To this end, we established a predictive protein panel consisting of NBL1, LTBP4, APOC4, and GHR. These proteins reflect the critical pathological processes elucidated above, namely active fibrogenesis and progressive metabolic dysfunction, thereby providing robust biological justification for their utility as prognostic biomarkers. In the test cohort of 22 MASLD cirrhotic patients the model demonstrated an AUC of 0.93 for predicting future decompensation in initially compensated patients. The model’s high specificity, correctly identifying all patients who remained compensated, minimizes false positives, a desirable property when aiming to avoid unnecessary interventions. Concurrently, by detecting most patients destined for decompensation (our ‘high-risk’ compensated group), patients whom the model classifies as high-risk for future decompensation might warrant a different therapeutic approach, as certain interventions typically well-tolerated by stable compensated individuals could be less suitable for this progressing subgroup. Conversely, those predicted to remain stably compensated could benefit from such interventions.

External validation in an independent cohort with alcohol-related cirrhosis (n=120) yielded an AUC of 0.78, supporting partial generalizability across aetiologies. However, the decrease in performance from the MASLD test set was likely due to several factors: (1) potential overfitting to the small training dataset (n=24), (2) biological differences between MASLD and alcohol-related cirrhosis, and (3) differences in cohort characteristics, such as prevalence of decompensation (50% in test vs. 77% in validation) and the progression profile of the test set (which was initially compensated and later decompensated). These factors likely contributed to an upward shift in model scores within the validation cohort. When the test-set optimal threshold of 0.49 was applied to the validation cohort, sensitivity remained high but specificity substantially decreased. Compared to the test set, this reflected an increase in false positives. Consequently, a higher cutoff was needed to effectively limit false positives in this cohort. The validation cohort’s Youden threshold of 0.70 provided more balanced performance and a higher LR+, demonstrating better diagnostic utility for this specific population. Threshold selection should reflect aetiology and clinical priorities: lower cutoffs for broad case-finding to minimize missed cases, and higher cutoffs to target truly high-risk patients and reduce unnecessary interventions.

Beyond cross-sectional discrimination, the model showed prognostic value in the validation cohort. Higher baseline probability scores were associated with shorter time to liver-related events (HR=1.32), and Kaplan–Meier curves demonstrated significant risk separation (log-rank p=0.0444). These findings suggest that the panel not only identifies current decompensation but may also capture future progression risk. Assessing the prognostic value of panel proteins, NBL1 showed significant association, whereas LTBP4, APOC4, and GHR did not show independent predictive value. This likely reflects the distinct aetiologies. Improved prognostication for alcohol-related cirrhosis could therefore benefit from aetiology-specific weighting, model recalibration, or inclusion of more disease-specific proteins. Despite a performance decline from the MASLD test cohort, the model’s validation was clinically meaningful: an AUC of 0.78 confirmed its diagnostic relevance, and an HR of 1.32, supporting its value for patient management.

## Conclusion

We developed a novel, informed 4-protein panel (NBL1, LTBP4, APOC4, GHR) that predicts future decompensated cirrhosis in MASLD. Grounded in key pathophysiological processes such as metabolic dysfunction and fibrogenesis, the panel demonstrated strong discriminative ability. Its performance in an external validation cohort further supports broad generalizability across aetiologies. While further prospective validation in larger, diverse cohorts is needed, this proteomic signature offers a promising strategy for the early identification of at-risk compensated cirrhotic patients, enabling timely interventions to prevent decompensation and improve clinical outcomes.

## Study Limitations and Future Directions

Our study has several limitations that warrant consideration. The primary limitation is the relatively small sample size of the training and test datasets, which, despite the use of regularization and event-per-variable ratio considerations, may contribute to the model overfitting this dataset. The limited size of the test dataset also yielded too few events for a statistically time-to-event analysis; consequently, survival analyses were restricted to the larger external validation cohort. While our external validation cohort was larger, the different aetiology (alcohol-related cirrhosis versus MASLD) meant direct biological comparisons were challenging. Furthermore, our study focused on a specific set of proteins, and future multiomic approaches could uncover additional predictive markers.

## Abbreviations

ANCOVA: analysis of covariance
ALDOB: fructose-bisphosphate aldolase B
ADH1B1: all-trans-retinol dehydrogenase
ALDH1A1: aldehyde dehydrogenase 1A1
BMP: bone morphometric proteins
COCOM-DO: Copenhagen Cohort of MASLD, Diabetes and Obesity
CST3: cystatin C
CNDP1: carnosine dipeptidase 1
nDIA: narrow-window data-independent acquisition
FIB-4: Fibrosis-4 Index
FDR: false discovery rate
GH: growth hormone
GHR: growth hormone receptor
HE: hepatic encephalopathy
HR: hazard ratio
IQR: interquartile range
IGF1: insulin-like growth factor 1
IGFBP1: insulin-like growth factor-binding protein 1
LTBP4: latent TGF-*β*–binding protein 4
LC-MS/MS: liquid chromatography-tandem mass spectrometry
LSM: liver stiffness measurement
LR: likelihood ratio
MASH: Metabolic dysfunction-associated steatohepatitis
MASLD: Metabolic dysfunction-Associated Steatotic Liver Disease
NPV: negative predictive value
NBL1: neuroblastoma suppressor of tumorigenicity 1
PPV: positive predictive value
PRO-DOC2019: PROspective cohort study of Disease and Outcomes in Cirrhosis
PCA: principal component analysis
SLC38A10: solute carrier family 38 member 10
T2D: type 2 diabetes
TWSG1: twisted gastrulation protein homolog 1
VSIG4: V-set and immunoglobulin domain-containing protein 4

## Declaration of generative AI and AI-assisted technologies in the manuscript preparation process

During the preparation of this work the author(s) used Gemini 2.5, ChatGPT-5, and Claude Sonnet 4.5 to refine and polish the text. After using this tool/service, the author(s) reviewed and edited the content as needed and take(s) full responsibility for the content of the published article.

## Authors Contributions

Concept and Design: MLNT, LLG. Acquisition of samples: MPW, MT, PN, EBR, LEH, SBK, AEJ, NK. Analysis, and Interpretation of data: MLNT, LLG. Drafting the Manuscript: MLNT, LLG. Critical Revision of the Manuscript for Important Intellectual Content: All authors.

## Data Availability Statements

The mass spectrometry proteomics data have been deposited to the ProteomeXchange Consortium (http://proteomecentral.proteomexchange.org) via the PRIDE [1] partner repository with the dataset identifier PXD075149 The python code with the data analysis is available in the GitHub repository at https://github.com/mlnth/COCO_decompensated_cirrhosis, released under MIT license.

## References

[1] Perez-Riverol Y, Bandla C, Kundu DJ, et al. “The PRIDE database at 20 years: 2025 update”. In: Nucleic Acids Research 53.D1 (2025), pp. 543–553.

[2] Younossi ZM, Golabi P, Paik JM, Henry A, Van Dongen C, and Henry L. “The global epidemiology of nonalcoholic fatty liver disease (NAFLD) and nonalcoholic steatohepatitis (NASH): a systematic review”. In: Hepatology 77.4 (2023), pp. 1335–1347.

[3] Tacke F, Horn P, Wai-Sun Wong V, et al. “EASL–EASD–EASO Clinical Practice Guidelines on the management of metabolic dysfunction-associated steatotic liver disease (MASLD)”. In: Journal of Hepatology 81.3 (2024), pp. 492–542.

[4] Mansour D and McPherson S. “Management of decompensated cirrhosis”. In: Clin Med (Lond) 18.2 (2018), pp. 60–65.

[5] Zipprich A, Garcia-Tsao G, Rogowski S, Fleig WE, Seufferlein T, and Dollinger MM. “Prognostic indicators of survival in patients with compensated and decompensated cirrhosis”. In: Journal of Hepatology 32.9 (2012), pp. 1407–1414.

[6] Smith AD, Zand KA, Florez E, et al. “Liver Surface Nodularity Score Allows Prediction of Cirrhosis Decompensation and Death”. In: Radiology 283.3 (2016).

[7] Thomson MJ, Lok AS, and Tapper EB. “Optimizing medication management for patients with cirrhosis: Evidence-based strategies and their outcomes”. In: Liver International 38.11 (2018), pp. 1882–1890.

[8] Werge MP, Grandt J, Thing M, et al. “Circulating and hepatic levels of growth differentiation factor 15 in patients with metabolic dysfunction-associated steatotic liver disease”. In: Hepatology Research 55.4 (2025), pp. 492–504.

[9] Benoit SW, Ciccia EA, and Devarajan P. “Cystatin C as a biomarker of chronic kidney disease: latest developments”. In: Expert Review of Molecular Diagnostics 20.10 (2020), pp. 1019–1026.

[10] Huang X, Li Y, Jiang L, and al. et. “Comprehensive pan-cancer investigation of carnosine dipeptidase 1 and its prospective prognostic significance in hepatocellular carcinoma”. In: Open Med (Wars) 19.1 (2024), pp. 2391–5463.

[11] Yang X, Ye X, Zhang L, Zhang X, and Shu P. “Disruption of LTBP4 Induced Activated TGFB1, Immunosuppression Signal and Promoted Pulmonary Metastasis in Hepatocellular Carcinoma”. In: OncoTargets and Therapy 13 (2020), pp. 7007–7017.

[12] Tsai TH, Song E, Zhu R, et al. “LC-MS/MS-based serum proteomics for identification of candidate biomarkers for hepatocellular carcinoma”. In: Proteomics 15.13 (2015), pp. 2369–2381.

[13] Wang W, Wang C, Xu H, and Gao Y. “Aldehyde Dehydrogenase, Liver Disease and Cancer”. In: International Journal of Biological Sciences 16.6 (2020), pp. 921–934.

[14] Privitera G, Spadaro L, Marchisello S, Fede G, and Purrello F. “Abnormalities of Lipoprotein Levels in Liver Cirrhosis: Clinical Relevance”. In: Digestive Diseases and Sciences 63.1 (2018), pp. 16–26.

[15] Borrego MC, Rio-Moreno M del, and Kineman R. “Towards Understanding the Direct and Indirect Actions of Growth Hormone in Controlling Hepatocyte Carbohydrate and Lipid Metabolism”. In: Cells 10 (2021), p. 2532.

[16] Yuen KCJ, Hjortebjerg R, Ganeshalingam AA, Clemmons DR, and Frystyk J. “Growth hormone/insulin-like growth factor I axis in health and disease states: an update on the role of intra-portal insulin”. In: Frontiers in Endocrinology 15 (2024), pp. 1664–2392.

[17] Ma IL and Stanley TL. “Growth hormone and nonalcoholic fatty liver disease”. In: Immunometabolism (Cobham) 5.3 (2023), p. 00030.

[18] Saeki C, Kanai T, Ueda K, et al. “Insulin-like growth factor 1 predicts decompensation and long-term prognosis in patients with compensated cirrhosis”. In: Frontiers in Medicine 10 (July 24, 2023), p. 1233928.

[19] Nolan K, Kattamuri C, Luedeke DM, et al. “Structure of neuroblastoma suppressor of tumorigenicity 1 (NBL1): insights for the functional variability across bone morphogenetic protein (BMP) antagonists”. In: Journal of Biological Chemistry 290.8 (2015), pp. 4759–4771.

[20] Malinauskas T, Moore G, Rudolf AF, et al. “Molecular mechanism of BMP signal control by Twisted gastrulation”. In: Nature Communications 15 (2024), p. 4976.

[21] Wan S, Liu X, Sun R, et al. “Activated hepatic stellate cell-derived Bmp-1 induces liver fibrosis via mediating hepatocyte epithelial-mesenchymal transition”. In: Cell Death and Disease 15 (2024), p. 41.

[22] Liu B, Cheng L, Gao H, et al. “The biology of VSIG4: Implications for the treatment of immune-mediated inflammatory diseases and cancer”. In: Cancer Letters 553 (2023), p. 215996.

